# Angle-dependence and auxin asymmetry: towards the formulation of a universal theory of root gravitropism

**DOI:** 10.1101/2022.11.19.517037

**Authors:** K. Sageman-Furnas, M. Del Bianco, J. Friml, C. Wolverton, S. Kepinski

## Abstract

Gravitropism has been conceptualized through three main theories: the starch-statolith model for sensing, the Cholodny-Went model of growth control via auxin asymmetry, and the law of angle-dependence for behavior, in which the magnitude of gravitropic response increases with displacement from the vertical. While experimental data show that the generation of both Cholodny-Went-type auxin gradients and angle-dependent behavior requires statolith sedimentation, a link between auxin asymmetry and angle-dependence has not been demonstrated. Here, we use large scale reorientation assays of Arabidopsis roots, epidermal length measurements, and confocal microscopy to quantify auxin distribution and PIN localization during graviresponse. We show angle-dependence in auxin asymmetry and growth response, even at low stimulation angles. As such, our work integrates the three theories sensing, signal transduction and behavior into a single unified model of gravitropism and provides an important framework for exploring major outstanding questions in the field.

## Results and Discussion

Plants have evolved the ability to sense their orientation within the gravity field and adapt their growth and development, enabling them to maximize the capture of resources above and below ground. This process of perception and response to gravity, known as gravitropism, has been conceptualized through three main theories: the starch-statolith theory for sensing, the Cholodny-Went model for signal transduction and growth control, and the law of angle-dependence for behavior. Together, in the root, they form a picture in which, following statolith sedimentation in gravity-sensing columella cells, PIN-FORMED (PIN) proteins re-localize to direct auxin to the lower side of the root, inhibiting cell elongation and thereby driving tropic curvature (Friml et al., 2002, Li, Gallei & Friml, 2022). Evidence from constant stimulus studies have shown that angle-dependence, where the magnitude of gravitropic response increases with angular displacement, relies on the presence of sedimenting statoliths (Mullen et al., 2000), thus connecting angle dependence with the starch statolith theory. Similarly, a model of root gravitropism based on auxin distribution showed that auxin asymmetry is lost in mutants lacking starch and, hence, robust statolith sedimentation (Band et al., 2012). However, the same study suggested that the graviresponse is uncoupled from auxin asymmetry at angles below 48° from the vertical (Band et al., 2012). Given the robust response to gravistimulation at angles well below 48° (Mullen et al., 2000), we sought to reevaluate the relationship between auxin asymmetry and angle dependent gravitropic response using newer tools for highly sensitive, more quantitative reporting of auxin gradients in graviresponding roots. We show that gravitropism in Arabidopsis is angle-dependent and governed by quantifiable auxin gradients from 30°, and provide a mechanistic framework towards the formulation of a universal theory of root gravitropism.

We first characterized in detail the gravitropic response of Arabidopsis with reorientation experiments starting at different angles (30°, 60°, 90°, 120°, 150°, and 170°; n > 76) (Figure 1A). Vertically grown 5-day-old Col-0 seedlings were gravistimulated in infrared light, and imaged at 30-minute intervals using a converted infrared camera with an 830 nm filter. To quantify the magnitude of gravitropic response in the root, we measured the average bend rate in the first hour after reorientation (Figure 1B). The initial bend rate gradually increased between 30° and 120° reorientation angles and then decreased between 150° and 170° (p < 0.05, 1-way ANOVA), with approximately 120° eliciting maximal response. These data are consistent with the seminal work of Sachs and more recent research on Arabidopsis and other species ((IIno, Tarui & Uematsu, Mullen et al., 2000, Galland, 2002, Chauvet et al., 2019). Next, we looked for angle-dependence in the biophysical growth response by quantifying the reduction in the length of the lower side of the root as a parameter for curvature (1C). Col-0 plants were gravistimulated at 0°, 45°, 90°, and 135° in the same set-up used for reorientation experiments. We chose these angles as being representative of a smaller angle, of the horizontal, and of an angle slightly above that inducing the fastest bend rate (Figure 1A). Four hours after reorientation, roots were carefully mounted on glass slides with propidium iodide and imaged on a Zeiss LSM 880 Axio Imager 2 confocal. This analysis showed that the decrease in the length of the lower side significantly increased with reorientation angle (p < 0.05, 1-way ANOVA).

**Figure 1.**
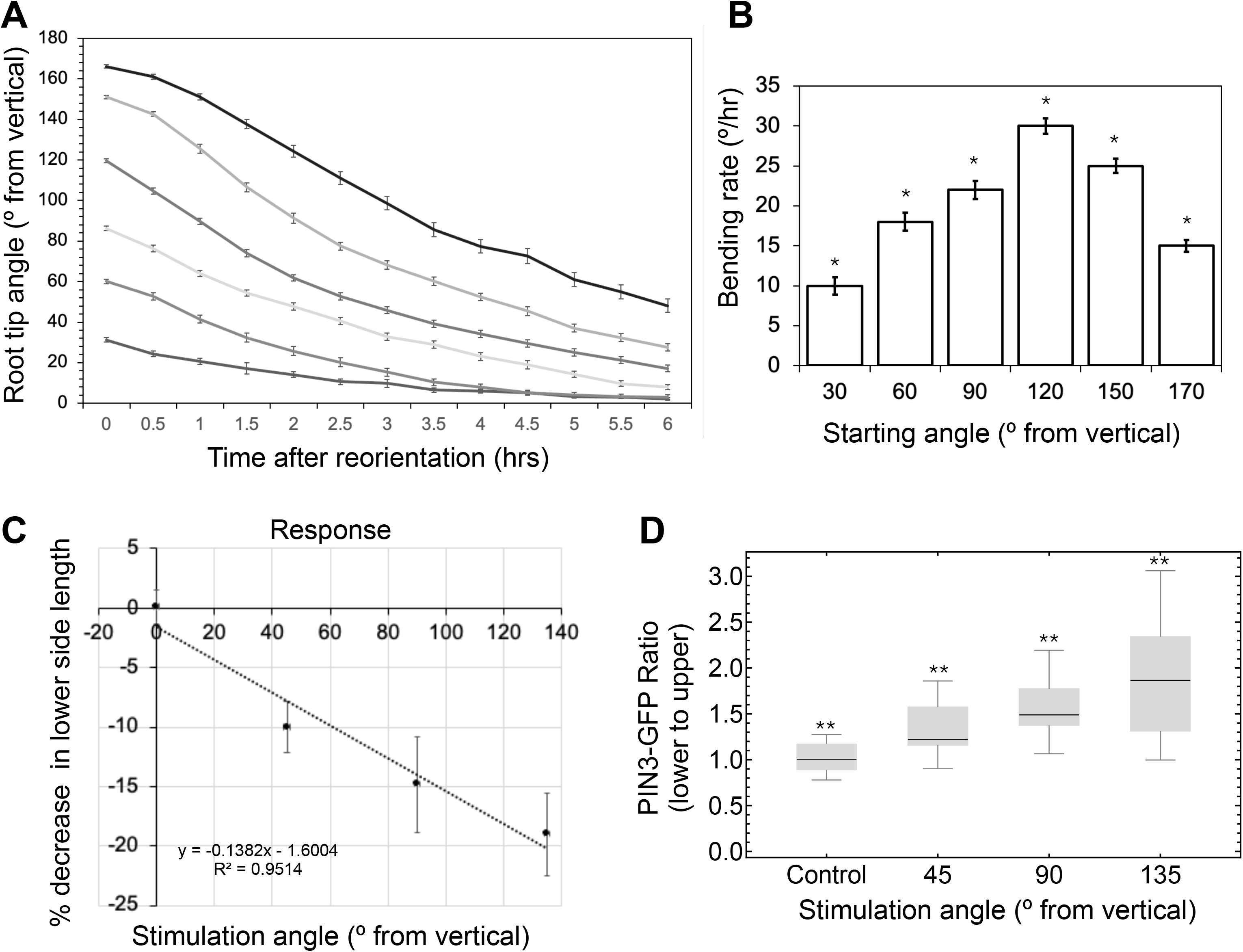
Gravitropic responses of Arabidopsis and PIN re-localization are angle-dependent. (A) Gravitropic response kinetics assays of Arabidopsis at different stimulation angles (30°-170°). (B) The first hour bending rate for each angle, n > 76 at each angle, bars represent s.e.m., stars represent p < 0.05, 1-way ANOVA with Tukey’s HSD test between all angle pairs. (C) Epidermal curvature of the root epidermis. Roots (5-7 each angle) were gravistimulated 4 hours in IR, allowed to acclimate for 1 hour and then imaged with a Zeiss LSM 880 Axio Imager 2 confocal (D) Ratio of lower to upper localization of PIN3::PIN3-GFP, n > 17 at each angle, stars represent p < 0.01, 1-way ANOVA with Tukey’s HSD test between all angle pairs. Box signifies the upper and lower quartiles and the median is represented by a black line within the box. The whiskers represent the highest and lowest values within a set.

To study the relationship between bending response and auxin gradients, we used the *R2D2* reporter, which, combined with vertical imaging, allows for quantitative inference of auxin levels (Liao et al., 2015). Following one hour of acclimation, five-day-old *R2D2* seedlings were imaged through the root mid-plane using a vertical-stage confocal microscope setup (Figure 2B). Roots were then gravistimulated at different angles (30°-120°) and imaged after 40 minutes. Relative auxin levels were calculated for each nucleus of the upper and lower root epidermis (Figure 2A-D). Since we found inherent differences in *R2D2* signal between hair and non-hair cells (Figure 2B, Student’s t-test, p < 0.01), we compared ratios between the same cell type (Figure 2C). Using this method, we found that quantifiable auxin gradients were present at 30° and that auxin gradients were correlated with stimulation angles between 30° and 120° (Figure 2D, R^2^ = 0.978). Other experiments with the newer *DR5v2* construct and a non-vertical stage approach also showed angle-dependent auxin gradients but failed to show a significant auxin gradient at 30° (Figure S1), highlighting the importance of a vertical stage approach and a more sensitive auxin reporter. These data suggest that auxin gradients direct gravitropic bending in Arabidopsis roots even at 30° and support the idea that angle-dependent auxin gradients dictate the magnitude of root gravitropic response.

**Figure 2.**
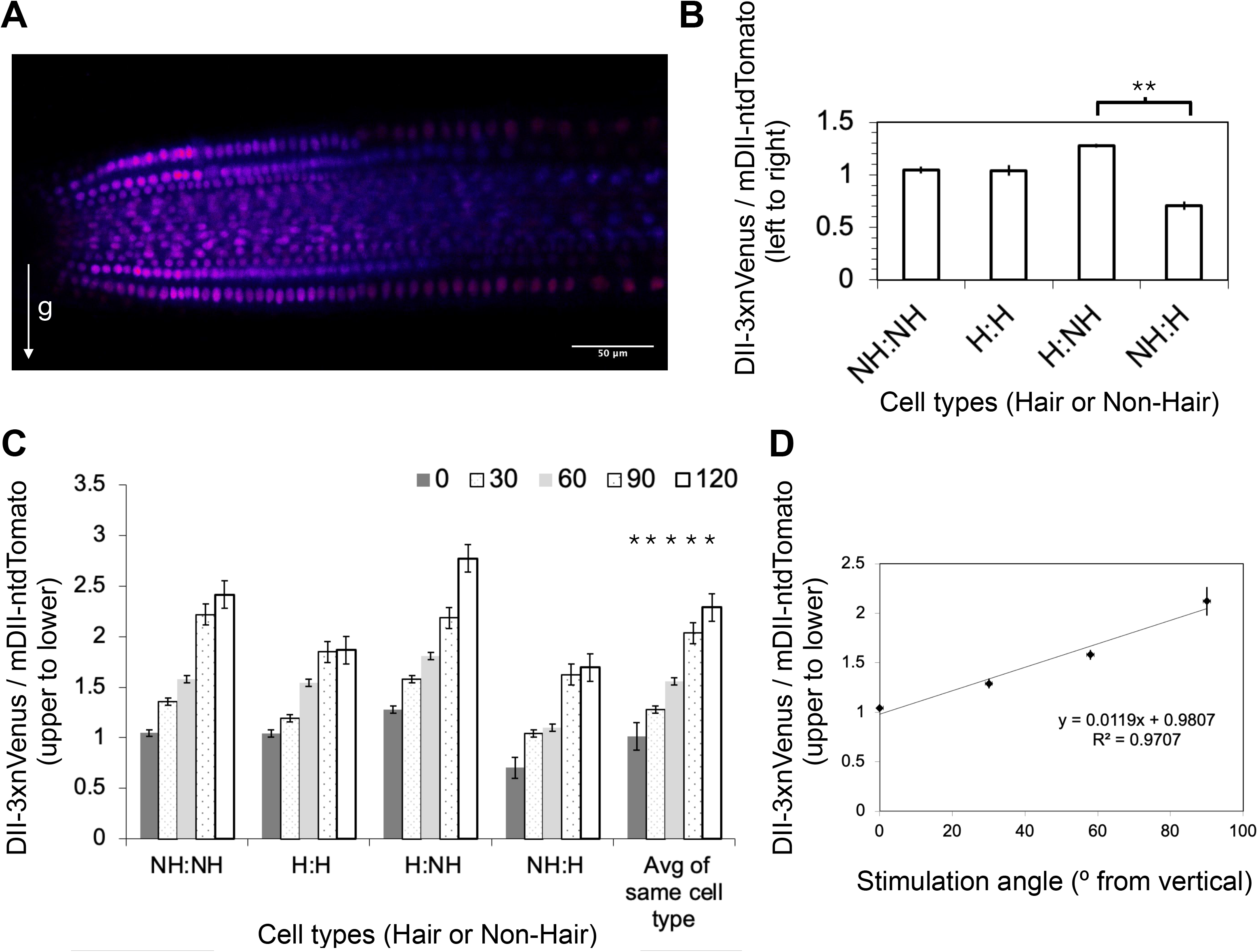
Angle-dependent auxin asymmetries in the Arabidopsis primary root tip can be quantified using *R2D2*. (A) An example image used for analysis of inferred auxin asymmetries using the *R2D2* reporter. (B) Venus and ntdTomato fluorescence of *R2D2* comparing auxin asymmetries between Hair and Non-Hair cell types in vertically growing roots, n > 6, stars represent p < 0.01, Student’s t-test. (C) Inferred auxin asymmetries of roots gravistimulated for 40 minutes at different angles. (D) Inferred auxin asymmetries of gravistimulated roots based on auxin measurements in only the same type of cells, n > 12 for each cell type and each angle, stars represent p < 0.05, 1-way ANOVA with Tukey’s HSD between all angle pairs when compared within the same cell type. All bars represent s.e.m.

Since PINs are known to mediate asymmetric auxin transport during graviresponse, we assessed for angle-dependent differences in the localization of PIN3 (Kleine-Vehn et al., 2010). *PIN3::PIN3-GFP* roots were imaged after a 40-minute gravistimulation at 0°, 45°, 90°, and 135°. As it is impossible to discern PINs localized to adjacent plasma membranes (PM), we used only the cells at the edge of the *PIN3* expression domain (Von Wangenheim et al., 2017). The upper vs. lower subcellular localization was calculated as a ratio between the mean PIN3-GFP fluorescence of the lower PM edge and of the upper PM edge of each root for each stimulation angle (Figure 1D). These data suggest that PIN3 is targeted to the lower side PM of the columella cells to a greater extent with increasing stimulation angle, indicating that angle dependence is already present as auxin transport asymmetry in the gravity-sensing cells.

Our work shows that the major components of root gravitropic response exhibit angle-dependence. Of particular note is demonstration of angle-dependence in Cholodny-Went-based auxin asymmetry and growth response that can be traced back to angle-dependent variation in PIN protein asymmetry in the gravity-sensing columella cells, even at low stimulation angles. More work will be needed to understand the proportionality between the different elements of the response and the presence of possible time-dependent features (Levernier, Pouliquen & Forterre, 2021). By integrating all of the separate theories for elements of root graviresponse, the work presented here represents an important step forward in our understanding of the biology of gravitropism and a framework for exploring major outstanding questions in the field.

---

**Figure S1.**
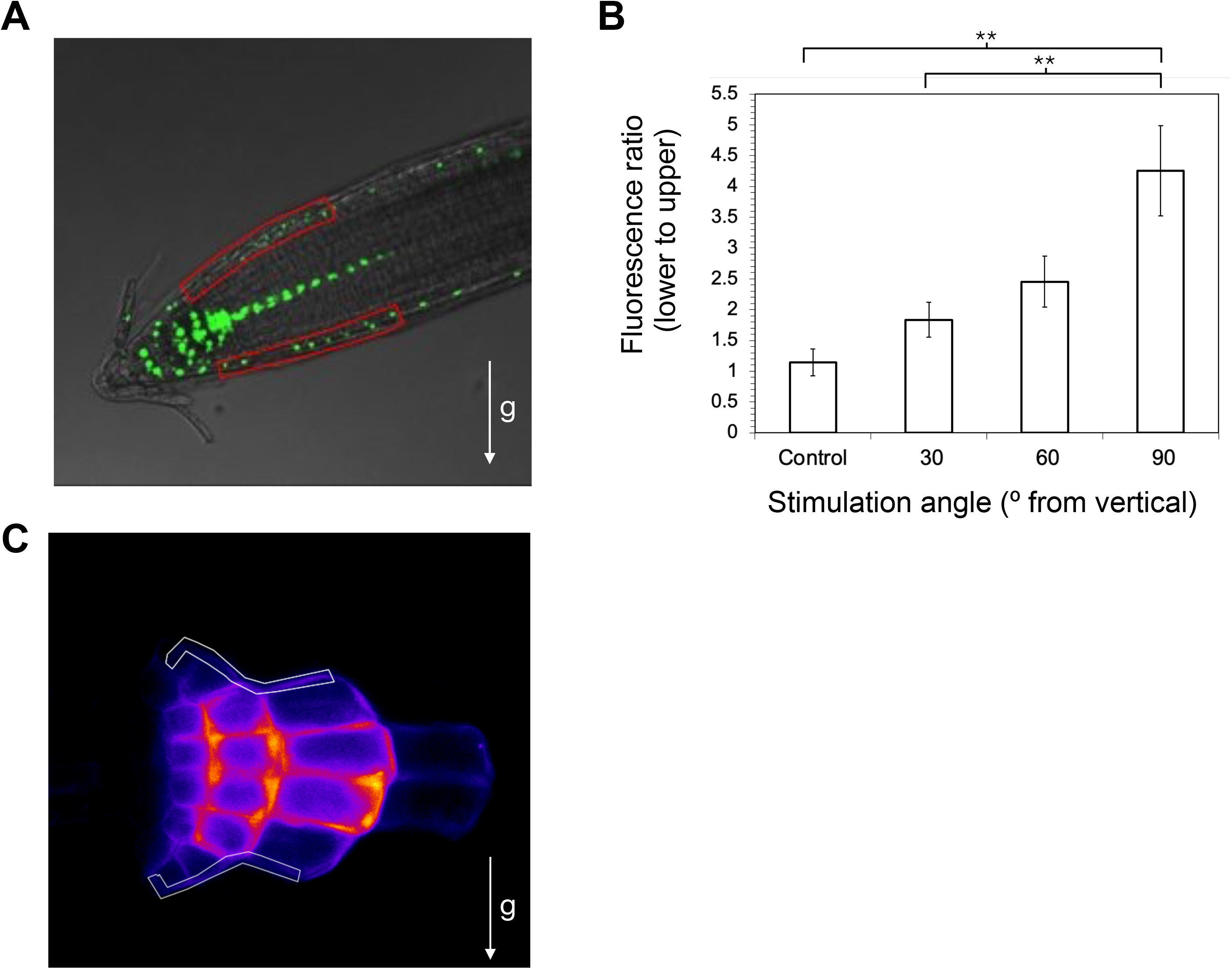
Angle-dependent auxin asymmetries visualized using *DR5v2*. (A) An example image used for analysis of inferred auxin asymmetries using the *DR5v2* reporter. Red boxes represent lower and upper portions. (B) Inferred auxin asymmetries stimulated at different angles using ROTATO (Mullen et al., 2000), n > 6 each angle, stars represent p < 0.01, 1-way ANOVA with Tukey’s HSD results showing significance between pairs. (C) Example image of a gravistimulated PIN3::PIN3-GFP root from Figure 1D.

## Notes

### Competing Interest Statement

The authors have declared no competing interest.

